# Preferential binding of human BRAC1 protein with open X-like conformation of Holliday junction, a homologous recombination intermediate

**DOI:** 10.1101/2024.08.23.609282

**Authors:** Sahil Hasan Kabir, K. Vishnupriya, Nibedita Pal

## Abstract

BRCA1 is a complex tumor suppressor protein involved in multiple critical cellular processes, e.g., DNA double strand break repair, cell cycle checkpoint, etc. BRCA1 depleted cells are reported to have decreased homologous recombination (HR) and promote error-prone non-homologous end joining for DNA damage repair. Holliday junction (HJ) is an important intermediate of HR. Although BRCA1 is shown to have a very high affinity for HJ and recruits several proteins at the DNA damage site, the question remains what the binding mode of BRCA1 protein with an HJ is. Using single-molecule Fluorescence Correlation Spectroscopy (FCS) we have shown that BRCA1 prefers an open X-like conformation of HJ and has a lesser affinity for stacked HJ. Further, using molecular docking and all-atom molecular dynamics simulation, we show that mostly charged and polar amino acids in the DNA binding region of BRCA1 form complex with HJ. Interestingly, most of those amino acids are reported to be places for missense changes.

## 1. Introduction

The integrity of the human genome is continuously compromised by cellular agents that modify the structure and function of DNA. DNA double-strand break (DSB) represents a particularly deleterious form of damage. Homologous recombination (HR) is an important pathway for DSB repair in mammalian cells.[1] Among several protein and protein complexes, breast cancer susceptibility gene 1 (BRCA1) protein is well characterized for its role in promoting DSB repair through HR.[2–4] BRCA1 mutated cancer cells are reported to be HR deficient and have an increased error-prone nonhomologous end-joining DNA repair mechanism.[5] This multifunctional protein comprises 1863 amino acids with a molecular weight of 220 kDa. The key features of the human BRCA1 protein are its ability to modulate protein-protein interaction via the N-terminal RING finger domain, two BRCA1 C-terminal (BRCT) domains, and proteinDNA interaction through the DNA binding region (DBR). The central region of the BRCA1 protein is largely disordered and is considered to play an important role as a flexible scaffold in its direct interaction with DNA and protein, thereby processing multiple signaling pathways.[6,7] Reports on the preferential binding of BRCA1 protein to non-canonical DNA structures propose the importance of its interaction with DNA in its regulatory role by selectively targeting non-B DNA structures such as four-way DNA Holliday Junction (HJ), G-quadruplex DNA, superhelical DNA, etc.[8,9] Its preference for branched DNA structures, which occurs due to DSB repair through HR, without sequence selectivity[9] and accumulation at the DNA damage site in a sequence-independent way, reaffirms its potential role in DNA damage repair.[10,11] However, its interaction mechanism at the molecular level with non-B DNA structures, such as HJ in DNA repair, is yet to be elucidated. The role of certain critical amino acids in this interaction and the potential preference of BRCA1 protein for a particular conformation of a cruciform DNA is not addressed.

Our study focuses on BRCA1’s affinity towards four-way HJ DNA among the non-B DNA structures. HJ DNA possesses a branch point and stems, an inherent structural motif of cruciform DNA. It is an important intermediate of DSB-repair through HR and replication. Attempts have been made to identify the DBR of BRCA1 by biochemical studies.[12–17] Residues 340-554 were reported to contain a DBR with a dissociation constant for four-way HJ of 3.3 μM for this domain. The authors speculated that multiple interactions within this domain might enhance BRCA1’s affinity for DNA despite the low DNA binding affinity measured.[12] Using single-molecule Fluorescence Correlation Spectroscopy (FCS), our study showed that the full-length BRCA1 protein has an affinity in nM^-1^ range for an HJ. In our investigation, we have shown, for the first time, that BRCA1 protein prefers an open X-like conformation of HJ over stacked conformation. It has an affinity of 5.8 χ10^7^ M^-1^ at low Mg^+2^ (0.5 mM) concentration at which the open conformer of HJ is populated. However, its affinity decreases with increasing Mg^+2^ concentration. At very high Mg^+2^ (50 mM) concentration, the binding of BRCA1 with HJ is almost negligible (3.0 χ10^6^ M^-1^). It is to be noted that at higher Mg^+2^ concentrations, HJ predominantly adopts stacked conformations. Previous investigations reported the strong binding of BRCA1 to branched DNA at low Mg concentrations; however, BRCA1’s affinity for HJ at higher Mg concentrations was ignored.[9] Addressing this is of paramount importance as HJ is a highly dynamic structure and samples three major conformations (two staked and one open-planer) in the presence of divalent cations.[18–20] Although BRCA1 does not exhibit any sequence selectivity for DNA binding, its preference for non-B DNA structures suggests its ability to identify different DNA conformations.

Further, we used computational methods to unravel the binding mode of BRCA1 with an HJ DNA. As a large portion of BRCA1 is disordered, we have modeled only DBR comprised of aa340-554, which is reportedly responsible for DNA binding.[12] We employed homology modeling, molecular docking, and all-atom molecular dynamics (MD) simulation to find the residues of BRCA1 responsible for its interaction with DNA. We have observed that mostly positively charged amino acids interact with a stacked HJ. The positively charged amino acids in the DBR form several binding pockets to interact with HJ. Interestingly, these residues are reportedly conserved across species and places for missense changes. It indicates the importance of the animo acids in mediating BRCA1 protein’s interaction with a damaged DNA and potential loss of interaction resulting from mutation.

## 2. Materials and Methods

### 2.1 Assembling the Holliday Junction

Fluorescently labeled and unlabeled Holliday Junction strands were procured from Integrated DNA Technologies and Eurofins Scientific1. These DNA strands were utilized as received. The as received strands were dissolved in HPLC water (Merck) to prepare 200 μM stock concentration for each strand (see Table 1). Specifically, strand 1 was tagged with Cyanine 3 (Cy3).

**Table1:**
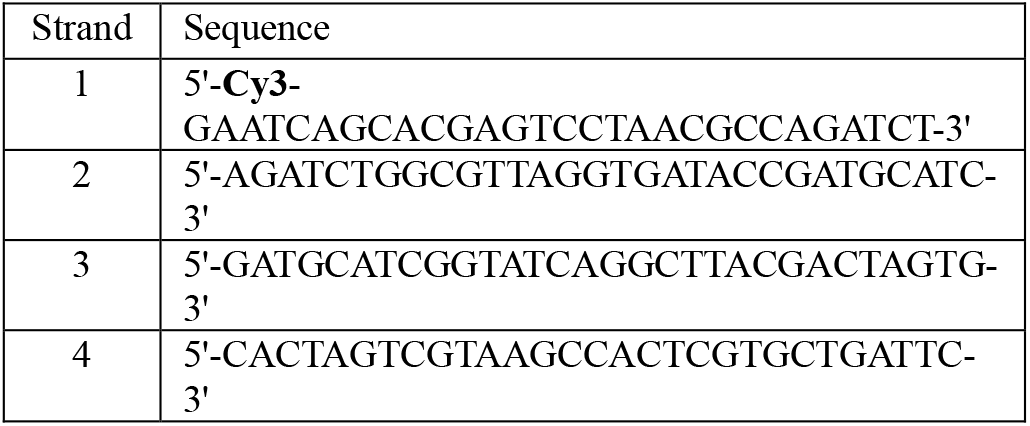
DNA sequences for annealing the Holliday junction.

Holliday Junctions were formed through annealing. The annealing process involved combining the four strands in a 1:1:1:1 molar ratio to achieve suitable DNA stock concentrations in MgCl_2_ (Spectrochem) and 20 mM Tris-HCl (Sigma-Aldrich) buffer at pH 7.5. This annealing was accomplished by heating the mixture to 90°C for 15 minutes and gradually cooling to 25°C over ∼5 hours. The annealed HJ was stored at 4°C for further use.

### 2.2 Protein

Recombinant Human BRCA1 protein was purchased from Abcam, UK (ab82204). BRCA1 was utilized after dissolving it in HPLC water (Merck) to produce a 480 nM stock.

### 2.3 FCS sample preparation

For FCS measurements, HJ concentration was kept constant at 2.5 nM, and BRCA1 concentration is gradually varied from 60 to 0 nM in the presence of 0.5, 1.0, 5.0, 12.5, and 50.0 mM MgCl_2_, 20 mM Tris-HCl, pH 7.5. After each dilution, the sample was incubated on ice for 15 min before FCS data acquisition. For each MgCl_2_ concentration, we had 11 samples with varying HJ:BRCA1 ratio, S1 = 1:24, S2 = 1:20, S3 = 1:15, S4 = 1:10, S5 = 1:8, S6 = 1:6, S7 = 1:4, S8 = 1:1, S9 = 1:0.5 and, S10 = 1:0.25, S11 = 1:0.

### 2.4 FCS data acquisition and analysis

FCS measurement involves correlating fluorescent fluctuations as a single fluorophore diffuses in and out of the confocal volume of ∼1fL. The duration for which this fluorophore or fluorophore-tagged molecule stays within the tiny volume depends on the diffusion coefficient. A typical model used to fit an autocorrelation curve for a molecule diffusing in 3D is given as

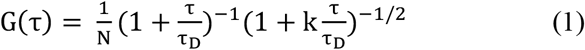

where, k is the square of the ratio of lateral (*r*) and axial (*l*) dimensions of the observation volume, τ_D_ is the is the average time a fluorescent molecule takes to diffuse through the observation volume. The diffusion coefficient (D) is related to τ_D_ by

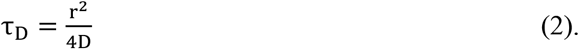

FCS measurements were carried out at room temperature in a home-built set-up built on an inverted confocal microscope. Using Rhodamine 6G as standard fluorophore (D = 4 × 10^−6^ cm^2^s^-1^), *r* is calculated to be 273 nm.[21] The average diffusion time of the Rhodamine 6G was maintained throughout all experiments.

In case of complex formation between two molecules (HJ and BRCA1 in our case), we used a two-population model with two different diffusion times for free and bound population of HJ (Eq. 3).[22]

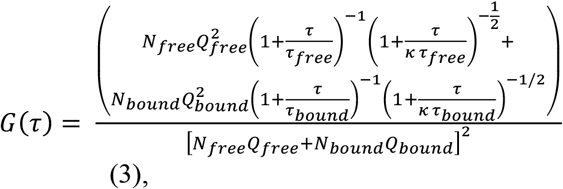

where, *N*_*free*_ and *N*_*bound*_ are the average number of free and bound species in the observation volume, respectively. *Q*_*free*_ and *Q*_*bound*_ are the brightness of the free and bound species and τ_free_ and τ_bound_ are their diffusion time, respectively. The values of *Q*_*free*_ and *Q*_*bound*_ are calculated from the ratio of photon counts and number of fluorescent molecules in the confocal volume at two extreme conditions (free HJ and completely bound HJ) and found to be 3.5 and 2.0, respectively. The correlation curves were fitted from 20.8 μs. This ensures only the diffusion-controlled part of the autocorrelation curve is considered. The molar fraction of the bound (χ_b_) and free (χ_f_) population are calculated as

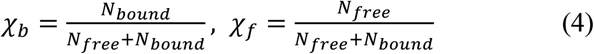

The association constant (*K*_*a*_) of BRCA1 to HJ can be calculated from *χ*_*b*_ from

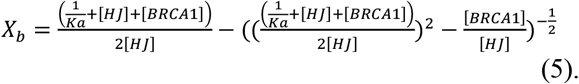

### 2.5 Molecular modeling and docking

The amino acid sequence of the human BRCA1 protein for residues 340-554 was retrieved from the Uniprot database with accession number P38398. We used the I-TASSER online server to predict the structure of this domain of BRCA1.[23] Each prediction in I-TASSER gives five models and based on the highest TM score and C score (Confidence score), we selected the best structure from each prediction (see Table S1 in *Supporting Information*). We employed I-TASSER thrice and selected three structures for further docking. The three predicted structures (model 1, 2 and 3) from I-TASSER were docked separately to a HJ (PDB:1DCW) using HADDOCK 2.4 web server.[24] We employed 5 solvation shells to establish a water layer between interacting partners.

### 2.6 Molecular Dynamics Simulation

The spontaneous association of DBR (aa 340-554) of BRCA1 with HJ (PDB:1DCW) was performed using Gromacs2022.2[25] at PARAM Smriti supercomputer. *Model2* was selected for this purpose. We placed the HJ at a distance ∼2.0 nm from the DBR at 4 orientations (see result section). All systems were solvated in a cubic water box using TIP3P water model. The systems were charge-neutralized, and the final NaCl salt concentration was 10 mM. System preparation was performed by AmberTools22 using AMBER ff19sb forcefield for proteins and BSC1 parameters for DNA HJ.[26–28] Simulations used periodic boundary conditions with the particle-mesh Ewald algorithm to handle electrostatic interactions, and a real space cutoff of 1.0 nm was set. Lennard-Jones interactions were truncated at 1.0 nm as well. All systems were subjected to 50,000 steps of the steepest descent minimization algorithm to remove the steric clash.[29] Further, the minimized systems were equilibrated into NVT and NPT phases for 5 ns each. The temperature (300K) and the pressure (1 bar) were maintained in all the systems using Vrescale, a modified Berendsen thermostat temperature coupling method, and the Parrinello–Rahman pressure coupling method as a barostat, respectively.[30,31] In all these steps, position restraint with a force constant of 1000kJ/mol/nm2 was applied on the DBR of BRCA1 and DNA HJ. Finally, we performed MD production of 100 ns for each system at NPT at 300K.

### 2.7 Analysis of molecular interactions

To assess the type of interactions and the type of amino acids participating in the interactions and overall picture of amino acid-nucleotide recognition, we investigate hydrogen bonds and van der Waals contacts. First, we selected the top two clusters from each docking based on the HADDOCK score to construct the datasets of hydrogen bonds and van der Waals contacts. The hydrogen bonds and van der Waals contacts in a complex were calculated using the Chimera visualization software with constraints relaxation of 0.6 Å and 60°.[32] Interacting residues are sorted according to their name, type, number of h-bonds, and interaction with the DNA backbone or the nitrogen bases. Based on the Haddock score, we have selected the best two clusters for further analysis for each model (see Table S2 in *Supporting Information*). Each cluster has 4 different complexes with possible binding modes and comparable HADDOCK scores. Thus, we have analyzed 8 complexes for H-bonds and VDW interactions for each model, 24 complexes in total. We listed the interacting amino acid residues in 8 complexes for each model and an overall distribution of the nature (polar, non-polar etc.) of the interacting residues is formed.

We also identified the van der Waals (VDW) interactions using ‘Find clashes/contacts’ module in Chimera. A VDW overlap cutoff of ≥ −0.4 Å was used. Like h-bonds, VDW contacts are computed between the protein and the HJ, and interacting residues are tabulated according to their name, type, etc.

In MD simulation, the changes in the minimum distance between DBR of BRCA and HJ, RMSF of the protein residues were calculated using the Gromacs module. Change in the number of H-bonds throughout the trajectory was calculated using the VMD Hbonds plug-in, where the donor-acceptor distance was set at 3.5 Å, and the angle cutoff was set at 30°. We also calculated the residue pair with the highest frequency of occurrence throughout the trajectory from the VMD Hbond plug-in.

## 3. Results and Discussion

### 3.1 FCS measurements

Auto-correlation curves of Cy3-labelled HJ in buffer and in presence of 0.5, 1.0, 5.0, 12.5 and 50.0 mM MgCl_2_ are shown in Figure 1a. All the auto correlation curves yielded a diffusion time of ∼190 μs. The rate of conformational exchange between *iso-I* and *iso-II* in HJ is reported to become faster with increasing salt concentration.[20,33] However, as we considered the part of autocorrelation functions only sensitive to diffusion, we cannot report any conformational dynamics happening in the timescale shorter than 20 μs. It is to be noted that the HJ used in our study does not undergo any branch migration. Thus, all the autocorrelation curves of HJs in the buffer are anticipated to yield a similar timescale of diffusion (190 μs in our case), irrespective of the salt concentration. We take this as the diffusion time of free HJs (*τ*_*free*_), i.e., unbound HJ for further analysis.

**Figure 1.**
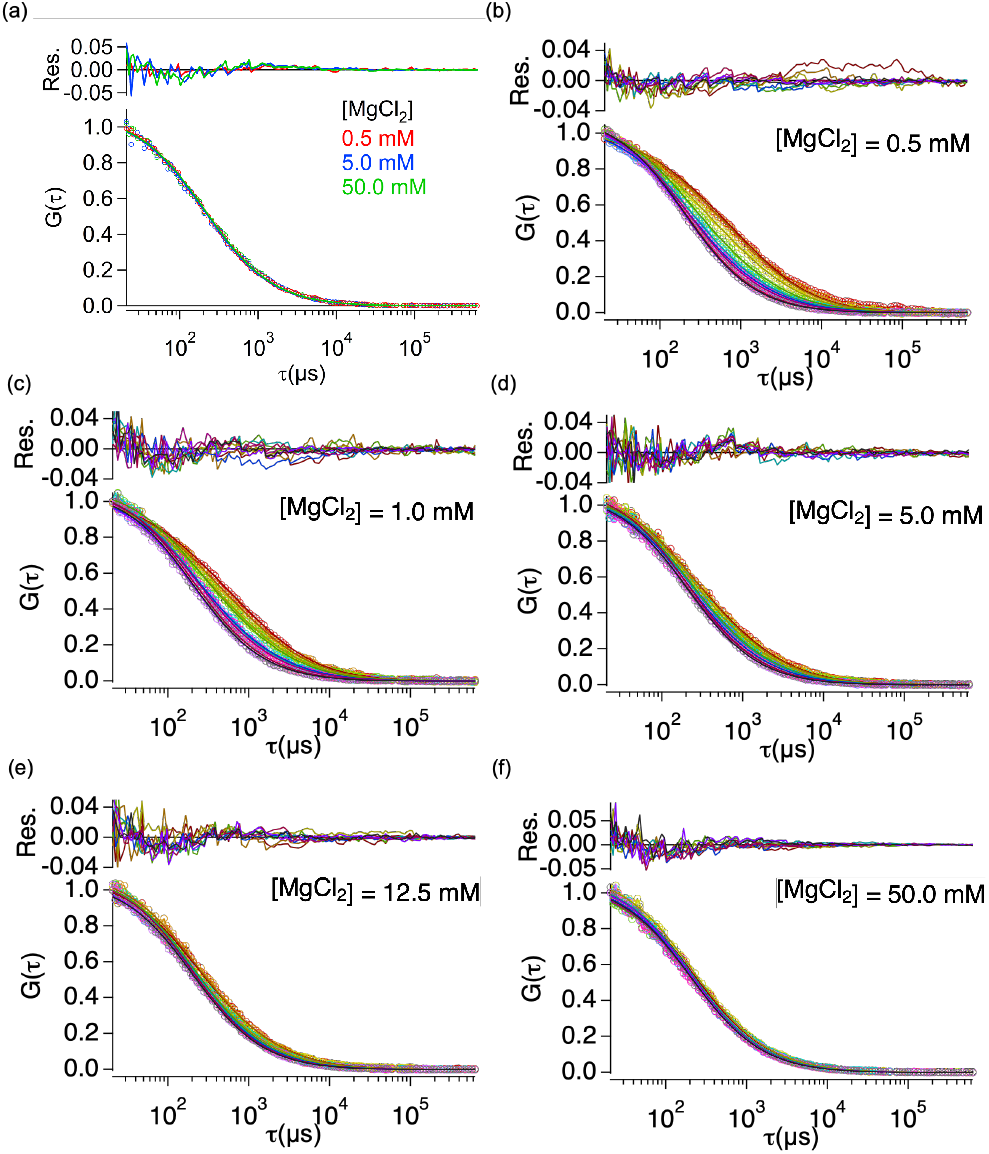
Auto-correlation curves of (a) HJ in buffer in presence of 0.5, 1.0, 5.0, 12.5, and 50.0 mM MgCl_2_ and that of HJ-BRCA1 in presence of (b) 0.5, (c) 1.0, (d) 5.0, (e) 12.5 and (f) 50 mM MgCl_2_ with varying BRCA1 concentrations. S1, S2, S3, S4, S5, S6, S7, S8, S9 and S10.

The binding of BRCA1 to HJ was followed by performing a reverse titration. HJ concentration was kept constant at 2.5 nM, and BRCA1 concentration gradually varied from 60 to 0 nM in the presence of 0.5, 1.0, 5.0, 12.5, and 50.0 mM MgCl_2_, 20 mM Tris-HCl, pH 7.5. After each dilution, the sample is incubated on ice for 15 min before FCS data acquisition. We have taken 2-3 data at each concentration to ensure the binding is complete before data acquisition. For each MgCl_2_ concentration, we had 11 samples with varying HJ:BRCA1 ratio, S1 = 1:24, S2 = 1:20, S3 = 1:15, S4 = 1:10, S5 = 1:8, S6 = 1:6, S7 = 1:4, S8 = 1:1, S9 = 1:0.5 and, S10 = 1:0.25, S11 = 1:0. At low MgCl_2_ of 0.5 mM, we except that HJ will be in an open X-like conformation due to high electrostatic repulsion between the arms. At higher BRCA1 concentrations, we observed that the autocorrelation curves are significantly slower and gradually become faster as BRCA1 concentration decreases (Figure 1b). This reflects the presence of slowly diffusing species at higher HJ:BRCA1 ratios resulting from the binding of BRCA1 with HJ. We performed a global fit analysis of the autocorrelation curves by eq 3, reflecting the coexistence of free and bound HJ species with individual diffusion time at any BRCA1 concentration. However, the fraction of bound and unbound HJ varies with BRCA1 concentration. Moreover, no bound population HJ is expected for S11. On the contrary, no free HJ species is expected at very high BRCA1 concentrations. As expected, we did not observe any shift in the autocorrelation curves for S1 and S2 at 0.5 mM MgCl_2_, suggesting that all the HJs are bound at these concentrations (Figure 1b). In the global fitting procedure, *k, Q*_*free*_, *Q*_*bound*_, τ_free_ and τ_bound_ are shared and *N*_*free*_ and *N*_*bound*_ are varied. The fractions of the bound and free populations are calculated using eq 4 and plotted in Figure 2a. As expected, χ_b_ increases with BRCA1 concentration as more and more complex is formed at saturates at HJ:BRCA1 = 1: 20 ratio (Figure 2a). We fitted χ_b_ with eq 5 which yielded the association constant *K*_*a*_ of 5.8 × 10^7^ M^-1^ (*K*_*d*_ = 17 nM). BRCA1 has been reported to exhibit extremely high affinity (*K*_*d*_ = 0.1-0.3 nM) for a flap DNA structure and reported to have comparable affinity for a branched DNA structure.[9] However, to the best of our knowledge, our study quantified the *K*_*d*_ of a full-length BRCA1 for a branched DNA HJ at the single molecule level for the first time. A higher *K*_*d*_ might be due to the differences in DNA arm length and Mg^2+^ concentration. We investigated the effect of MgCl_2_ on BRCA1 protein’s interaction with the same HJ. It is to be noted that at high Mg^2+^ concentration, HJ adopts a stacked conformation. At 1 mM MgCl_2_, *K*_*a*_ decreased almost twice (3.4 × 10^7^ M^-1^). At intermediate MgCl_2_ concentrations, 5.0 and 12.5 mM, *K*_*a*_ was measured to be 2.6 × 10^7^ M^-1^ and 12.5 × 10^6^ M^-^, respectively. At a much higher concentration, 50 mM MgCl_2_, negligible binding was observed (*K*_*a*_ = 3.0 × 10^6^ M^-1^). The differential binding of BRCA1 with HJ at different MgCl_2_ concentrations was evident from the FCS autocorrelation curves. Figure 3 shows the autocorrelation curves of S1, S2, S9 and S10 at 0.5, 1.0, 5.0, 12.5 and 50 mM MgCl_2_ (See *Supporting Information Fig S1* for the rest). The dependence on ionic conditions is evident at higher BRCA1 concentrations (S1, S2). However, at lower BRCA1 concentrations (S9, S10), there is little to no change among the autocorrelations curve with ionic conditions. Additionally, the only HJ in buffer did not exhibit any MgCl_2_ concentration dependency (Figure 1a). In Fig S2 we showed the normalized χ_b_ with MgCl_2_ concentrations. χ_b_ decreases rapidly with increasing MgCl_2_ concentration in for higher BRCA1 concentrations. However, at lower BRCA1 concentrations, the χ_b_ remains almost similar throughout the entire MgCl_2_ range. It should be noted that the conformational population of HJ depends on Mg^2+^ concentration. At low MgCl_2_ concentration, HJ prefers to stay in an open X-like conformation due to high electrostatic repulsion between the arms, whereas at high MgCl_2_ concentration, stacked conformations are populated. Thus, we infer that BRCA1 adopts a conformation selection while binding to HJ. HJ binding proteins, e.g., resolvase and DNA packaging proteins are reported to recognize stacked conformation of HJ and distort it to an open X-like conformation. Our finding of BRCA1 protein preferentially binding to an open X-like conformation may hold significance in DNA damage repair through HR. It would be worth investigating if such complex potentially facilitates the binding of other proteins essential for DSB repair.

**Figure 2.**
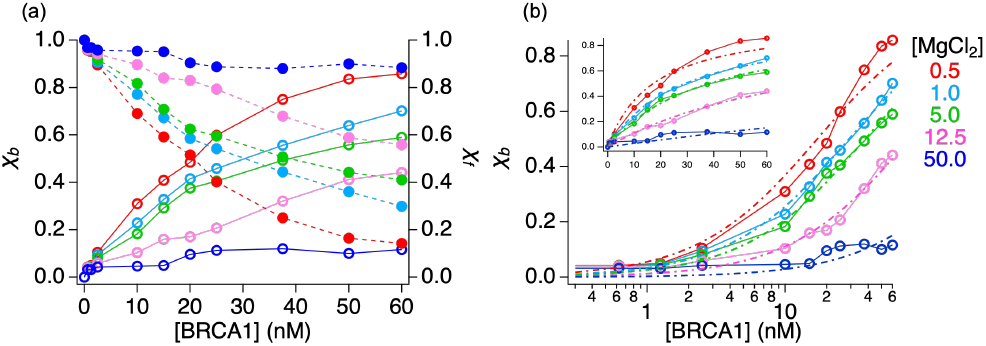
(a) Change in fraction of free (χ_f_, solid circle) and bound (χ_b_, open circle) HJ with BRCA1 concentration. (b) χ_b_ vs [BRCA1] in semilog plot with the fits by Eq 5. Inset, χ_b_ vs [BRCA1] in linear scale.

**Figure 3.**
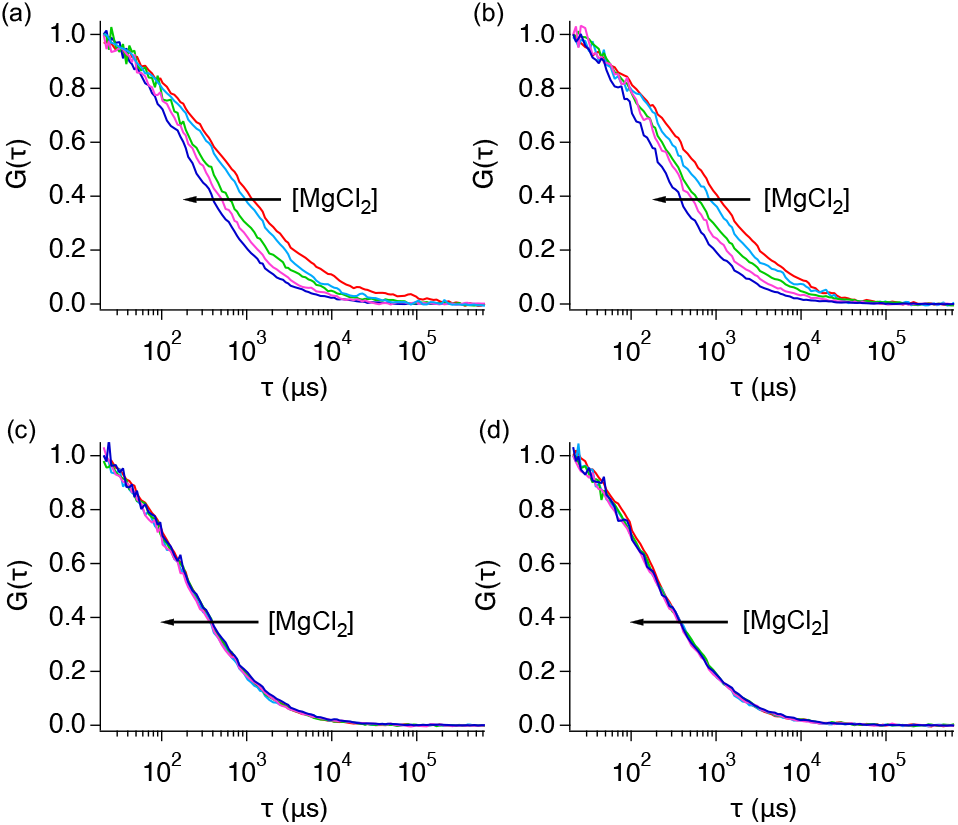
Autocorrelation curves of (a) HJ:BRCA1 = 1:24 (S1), (b)) HJ:BRCA1 = 1:20 (S2), (c) HJ:BRCA1 = 1:0.5 (S9), (d) HJ:BRCA1 = 1:0.25 (S10) with 0.5, 1.0, 5.0, 12.5 and 50 mM MgCl_2_.

### 3.2 Molecular modeling and docking

We leveraged molecular modeling and docking to find the amino acids that contribute most to the binding of the BRCA1 protein with HJ. We used I-TISSER for structure prediction.[23] A combined proteolysis, cloning, and spectroscopic study identified residues 340-554 of BRCA1 as minimal DBR.[12] Thus, we have selected this region for modeling. The DBR has high number of polar residues (66%, charged and uncharged) (see *Supporting Information Figure S3*). Based on various scores, we have selected three models (see *Supporting Information Table S1*). However, two of the structures were found to be identical.

Figure 4 shows the two structures predicted by I-TASSER. Corroborating with the previous reports, this region is mostly unstructured with short α-helices. A positively charged patch, rich in Lysine and other +vely charged amino acid residues, is found on the surface in all models (Figure 4c & d). One of the models indicated the presence of β-strands corroborating the previous report.[12] The occurrence of a significant number of loops was expected due to the presence of disorder-promoting residues, e.g., Alamine, Proline, Serine, Glutamate, Glutamine, Aspartate, etc. The three structures will be referred to as *model1, model2*, and *model3* from here onwards.

**Figure 4.**
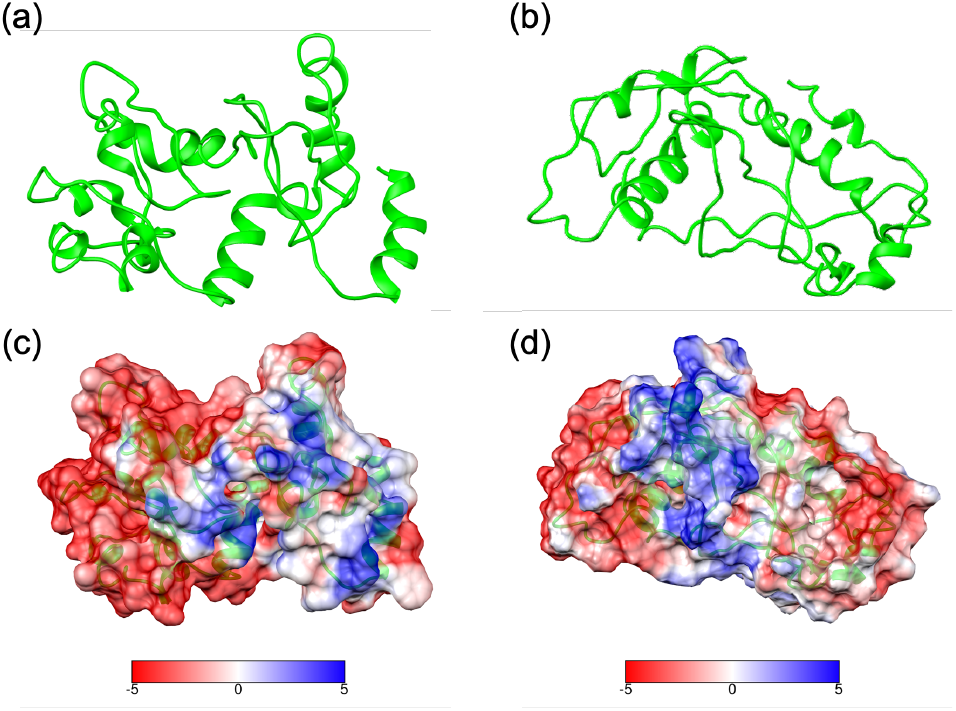
I-TASSER predicted secondary structures of DBR (aa340-554) of BRAC1: (a) Model1, (b) Model2 & 3. Calculated electrostatic surface of (c) Model1, (d) Model2 & 3.

Next, we docked the three predicted structures on a HJ. No crystal structure of an open X-like conformation of HJ is available. Thus, we docked 340-554 DBR on a stacked HJ (PDB: 1DCW). Representative docked complexes for each model are shown in Figure S4 in *Supporting Information*. In *model1* and *model3*, the modes of interactions are very similar in both clusters. However, in *model2*, the HJ interacts with the DBR of BRCA1 in two opposite sides in two models (*Supporting Information* Figure S4(b)). This might suggest that DBR of BRCA1 recognizes DNA structure, not DNA sequence. Although docking was performed with 5 solvation shells, no interfacial water molecules were found between HJ and BRCA1 protein in the docked structures. Mostly positively charged residues Lys and Arg were found to interact with the HJ. Polar uncharged Asn and Gln were also identified to interact with DNA. These residues mostly interacted with the DNA phosphate backbone through hydrogen bonds. The H-bonds in three representatives docked structures of *model1, model2* and *model3* are shown in Figure 5. Figure 6a-c shows the distribution of amino acid residues participating in the H-bond in three dockings. These amino acids form basic binding pockets in all three dockings to accommodate the negatively charged DNA backbone (Figure 6d-f). Overall, the negatively charged amino acids contribute significantly to the H-bonds regardless of the model and docking. This is typical for DNA-protein interaction driven by electrostatic interaction. Interestingly, the negatively charged amino acids, mostly Glu, contributed ∼5-10% of the H-bonds. These amino acids are found to interact directly with the nitrogen base of the DNA. It is well known that H-bonds with DNA sugar-phosphate backbones typically do not contribute to specificity to protein binding, but the two-thirds representation in the H-bond in our system emphasizes their significance in DNA structure recognition in a sequence-independent manner. We observed only ∼30-40% of the total H-bonds are directly between the polar amino acids and the nitrogen bases (see Figure 7a for a representative picture). Interestingly, the interacting nucleotides are mostly from the junction region of HJ (Figure 7b). However, understanding the sequence specificity of interaction requires more investigation in the future by changing the junction sequence.

**Figure 5.**
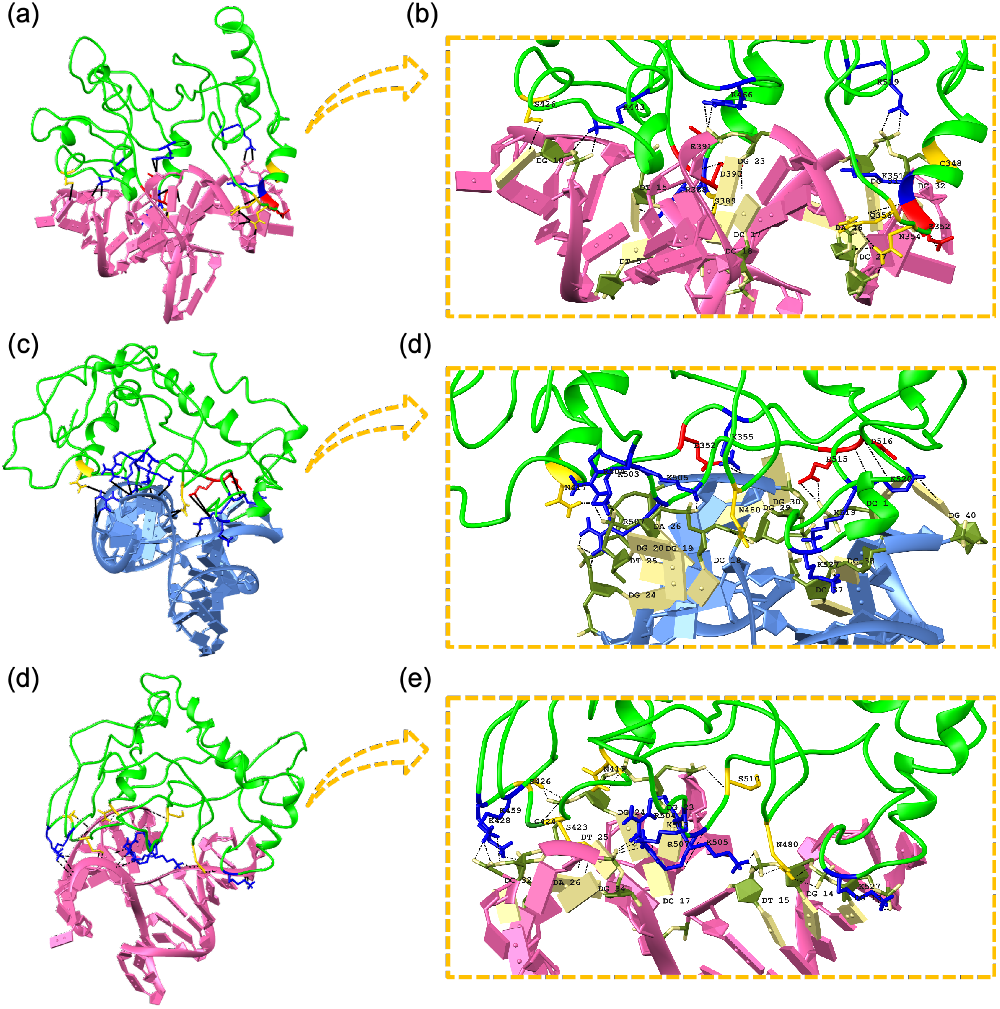
H-bonds between HJ and DBR in (a) model1, (b) zoomed-in picture of model1, (c) model2, (d) zoomed-in picture, (d) model3 and (e) zoomed-in picture. Polar uncharged residues are in yellow, polar positively charged residues are in blue. and polar negatively charged residues are in red color.

**Figure 6.**
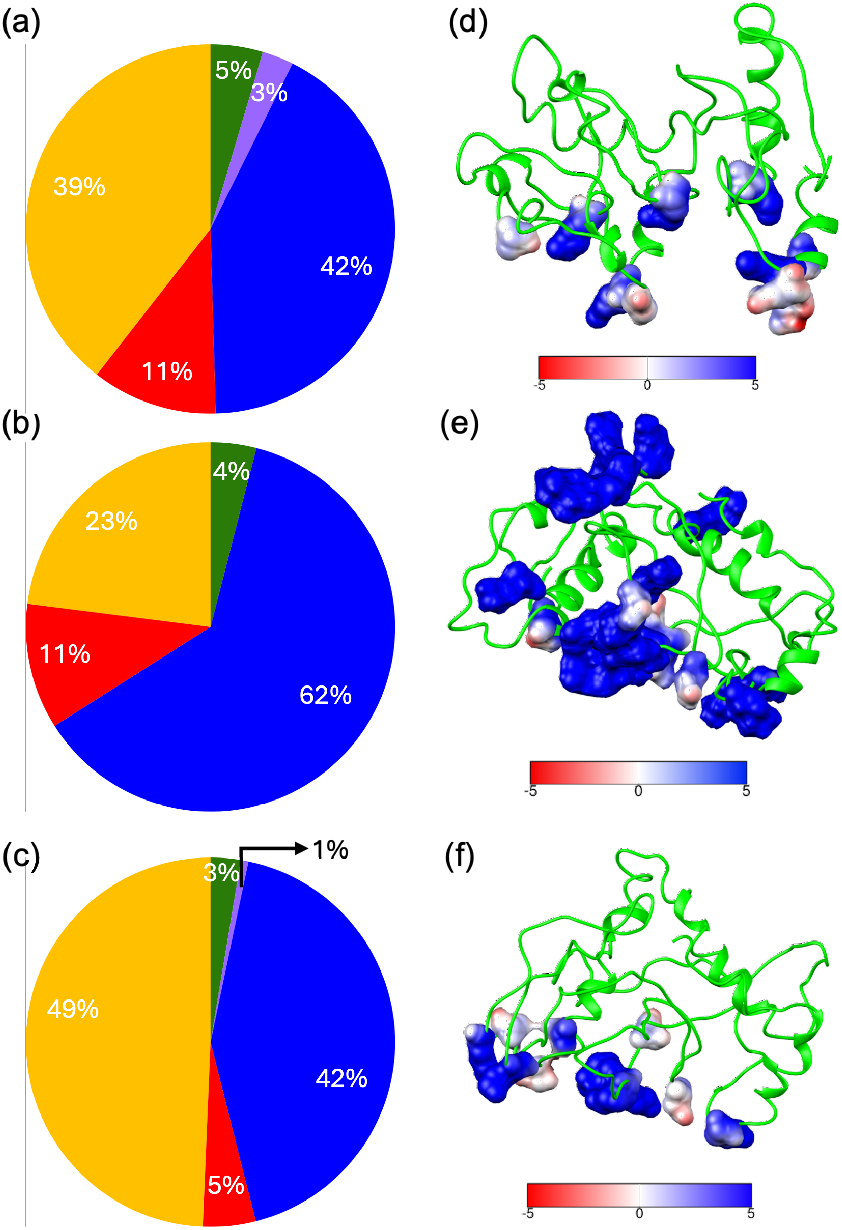
Distribution of the H-bond forming amino acid residues in docked structures of (a) model1, (b) model2 and model3. Polar uncharged residues are in yellow, polar positively charged residues are in blue, nonpolar aliphatic residues are in green, aromatic residues are in purple and polar negatively charged residues are in red color. Coulombic surface of the positively charged and polar uncharged residues combined in docked structure of (d) model1, (e) model2 and (f) model3. Coulombic surface coloring defaults: ε = 4r, threshold ± 5 kcal mol^-1^e^1^.

**Figure 7.**
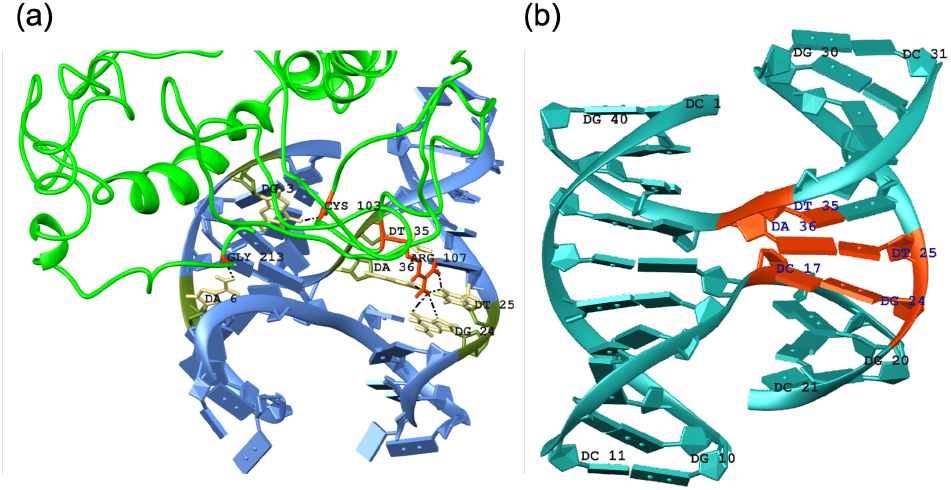
(a) H-bonds between amino acids nitrogen bases of the junction, (b) Nucleotides at the junction (orange) directly participating in H-bonds with DBD of BRCA1.

We observed higher VDW interactions between the DBR and HJ than the number of H-bonds. (Table S3 in Supporting Information). As anticipated, we observed mostly polar residues, e.g., ASP, GLN, SER, THR, ARG, LYS, GLU and ASP, contributed in VDW interaction (Figure 8).

**Figure 8.**
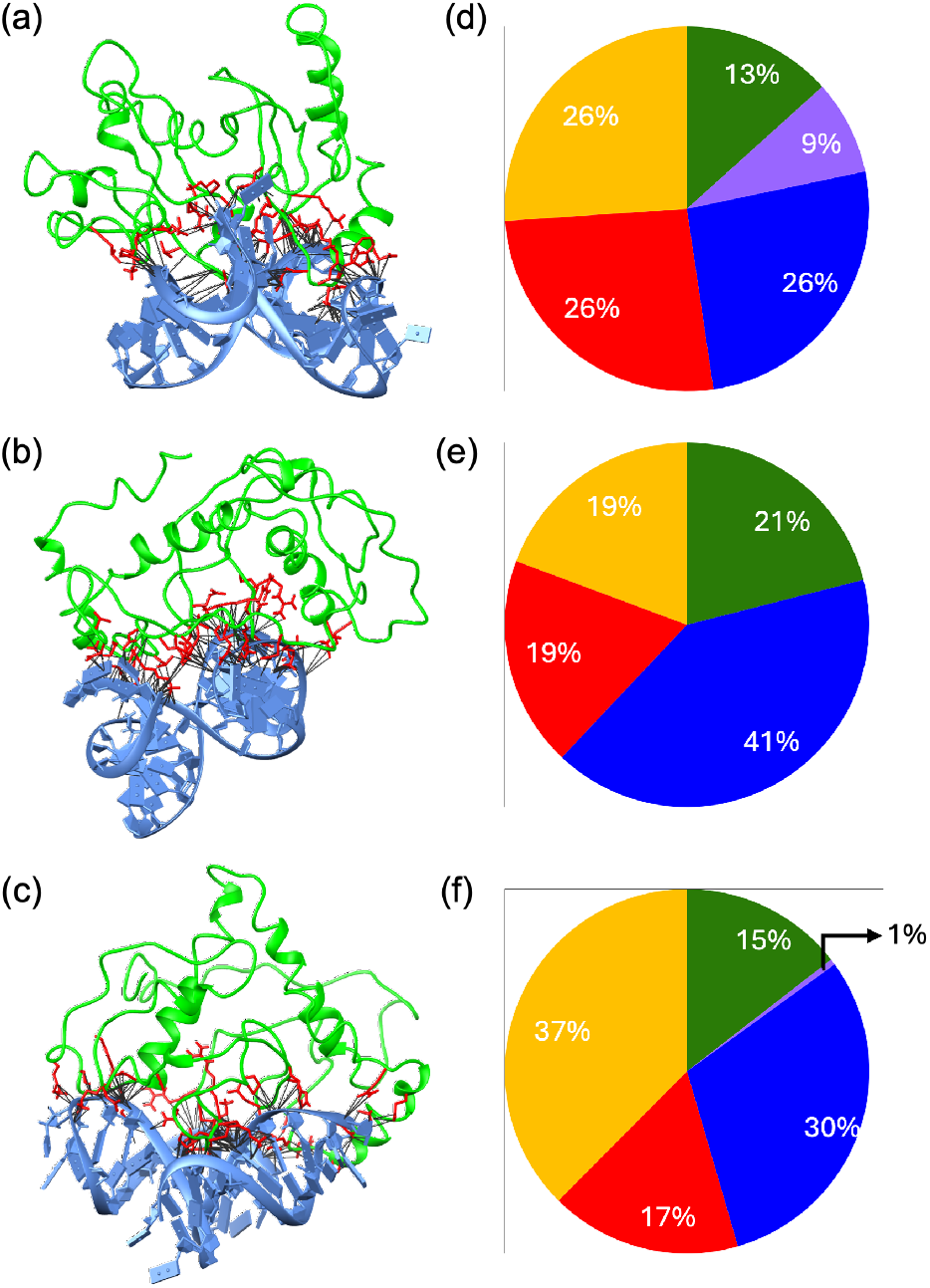
Representative VDW contacts in docked complexes of HJ with (a) model1, (b) model2 and (c) model3. (d-f) Distribution of the corresponding interacting amino acid residues. Polar uncharged residues are in yellow, polar positively charged residues are in blue, nonpolar aliphatic residues are in green, aromatic residues are in purple and polar negatively charged residues are in red color.

### 3.3 Molecular Dynamics Simulation

We investigated the spontaneous interaction of DBD of BRCA1 with HJ by employing all-atom molecular dynamics simulation. We performed four independent 100 ns simulations where DBR is placed at ∼2 nm from HJ at four different orientations (see *Supporting Information* Figure S5). This ensured a diverse sampling of binding modes. Previously, such an approach was adopted to provide an interaction model of the intrinsically disordered part of BRCA1 with a duplex DNA.[34] The study revealed a head-on interaction between the IRD region of BRAC1 and DNA duplex-end, suggesting its role in DNA damage repair. We observed spontaneous binding (defined as intermolecular distance <0.4 nm) between DBR and HJ within the first 20 ns in three out of four initial configurations. Once the complex was formed, it remained stable for the rest of the simulation (Figure 9). The fast binding suggested a strong propensity of BDB of BRCA1 with HJ. The overall root mean square deviation (RMSD) of the DBD increased in the first 15-20 ns. Once the encounters happened, the RMSD remained stable (*Supporting Information* Figure S6(a)). Higher fluctuation in the protein structure is also reported for other intrinsically disordered protein due high plasticity.[35] Interestingly, the DBR of BRCA1 prefers binding sidewise to the HJ in two of the three encounters (Figure 10a). This corroborates the previous reports on BRCA1 protein’s affinity for a branched DNA structure. Although, in *position 4* one of the arms bends to facilitate interaction with DBR of BRCA1, overall, HJ showed much lower RMSD (*Supporting Information* Figure S(b)). Further, we calculated the H-bonds with time for all the positions. Except for position 2, in all other positions, the number of H-bonds increased to around 20 ns and then stabilized (Figure 10b). Figure 10c shows the H-bond occupancy of the residues over 100 ns for *positions 1, 3* and *4*. Positively charged residues like Lys 428, 450, 463, 467, 501, 503, 505, 519, 527; Arg 466, 496, 504, and polar uncharged residue Asn 455, 480, Gln 526, 541 contributed most to the interaction.

**Figure 9.**
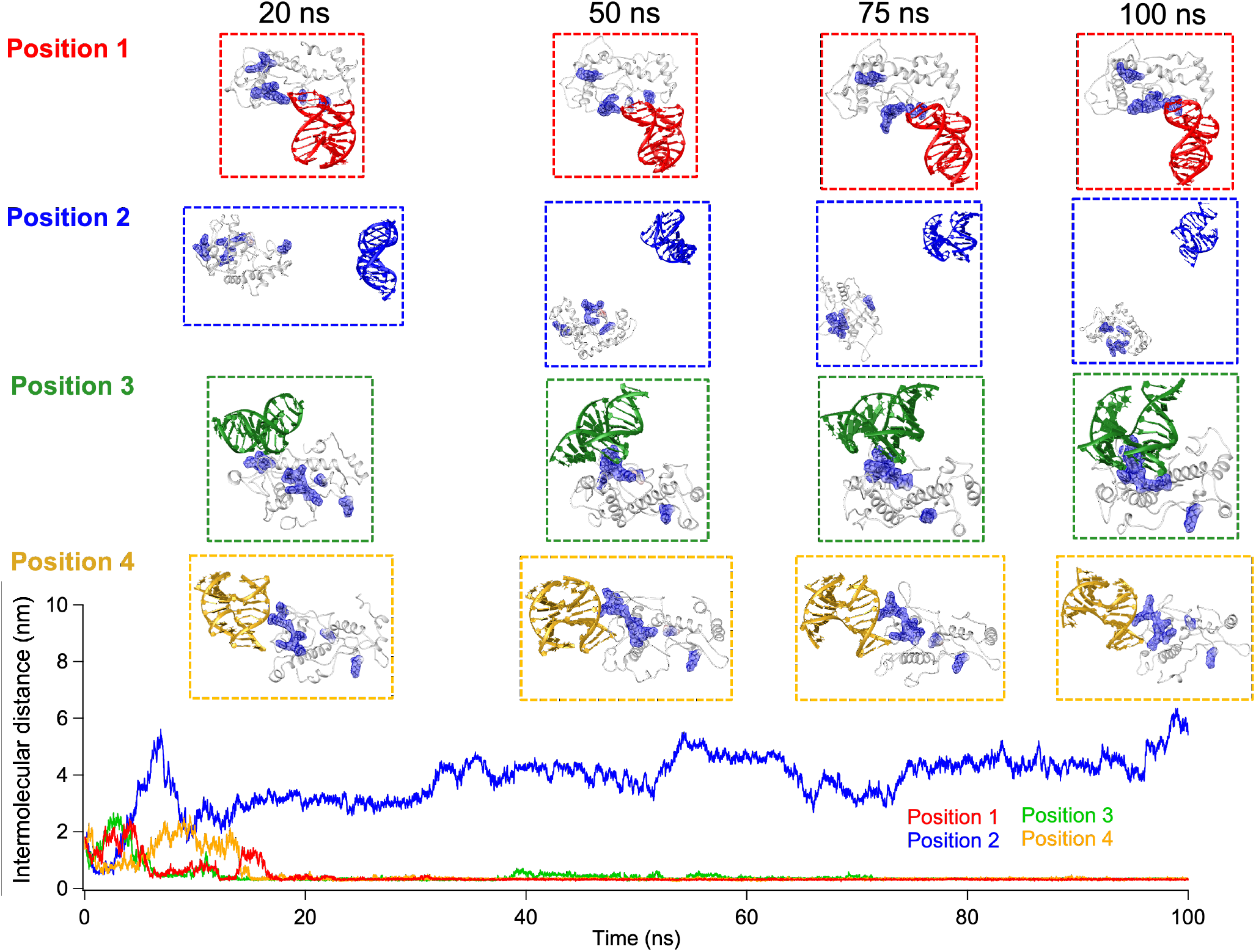
Upper panel – snapshots of DBD domain of BRCA1-HJ complex at 20 ns, 50 ns, 75 ns and 100 ns, Lower panel - Change in minimum distance between DBD domain of BRCA1and HJ with time.

**Figure 10.**
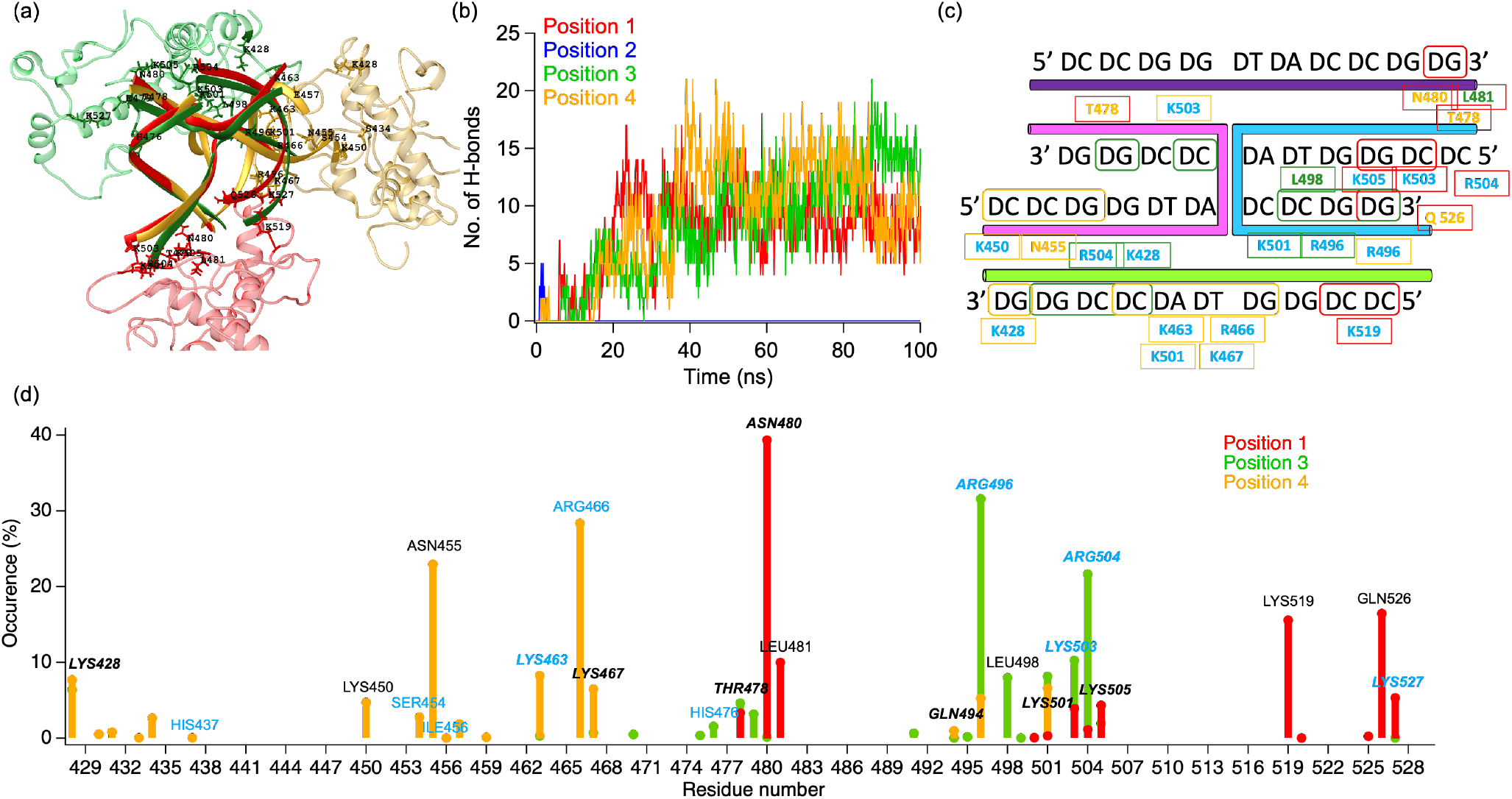
(a) Mode of interaction between DBR of BRCA1 and HJ, (b) Change in the number of H-bonds with time for all four simulations, (c) DBR binding sites on HJ and interacting residues, (d) occupancy of the interacting residues form DBR of BRCA1 protein. The amino acids found to be interacting with HJ both in docking and MD simulation are in bold and italics. The amino acid residues in light blue are places of missense changes in human BRCA1.[36] Color code: Red: Position1, Blue: Position 2, Green: Position 3 and Orange: Position 4.

Among these, Lys 467, 501, 503, 519, 526, Arg 466, 504, and Asn 480 were also found to form H-bonds in docking (Figure 5) as well. This establishes the important role these amino acids might play in stabilizing the HJ-DBD of the BRCA1 complex by forming a binding pocket. Interestingly, many missense changes in amino acid residues in this DBD are reported in the Breast Cancer Information Core (BIC) database. The interacting residues His 437, Ser 454, Ile 456, Arg 466,

His 476, Arg 496, Lys 463, 503, Arg 504, and Lys 527 in our study are reported for missense changes in BIC and other data databases (brcaexchange.org). Most of these residues are also reported to be strongly conserved across species, and these changes are predicted to affect the protein’s function.[36] This emphasizes the importance of these amino acids in BRCA1 protein’s interaction with DNA. The revelation that these amino acids are directly involved in the interaction with HJ likely explains why these places of missense mutations are essential for BRCA1’s function. The potential disruption of DNA binding activity due to mutations at these places may lead to genome instability and tumorigenesis.

Resembling the finding in molecular docking, we observed large number of VDW contacts between the DBR and HJ in MD simulation. As anticipated, we observed mostly polar residues contributed in VDW interaction (Figure 11).

**Figure 11.**
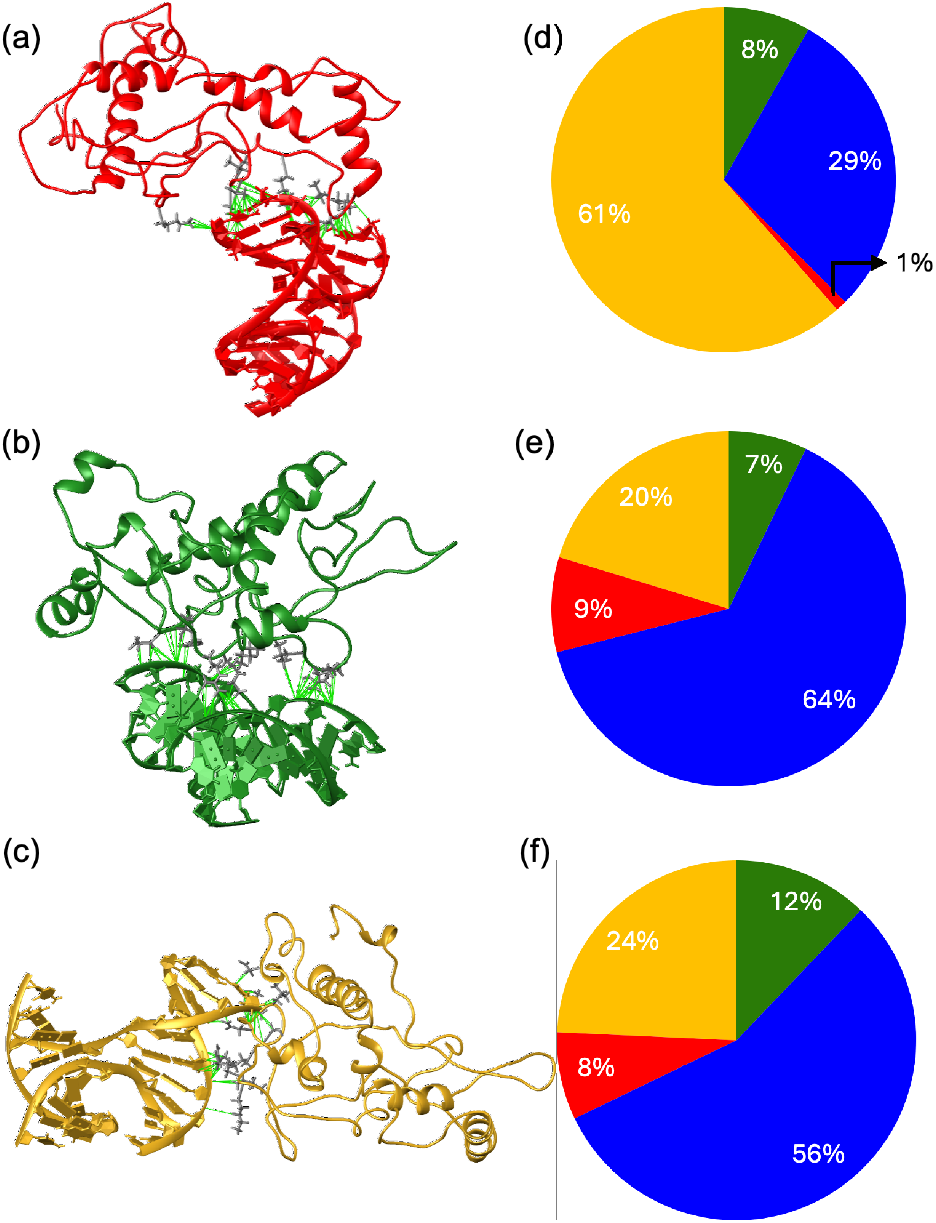
Representative VDW contacts in spontaneously formed complexes of HJ with model2 in (a) position 1, (b) position 2, and (c) position 3. (d-f) Distribution of the corresponding interacting amino acid residues. Polar uncharged residues are in yellow, polar positively charged residues are in blue, nonpolar aliphatic residues are in green and polar negatively charged residues are in red color.

## 4. Conclusion

Mutations in the BRCA1 gene predispose women to a very high risk of breast and ovarian cancer. The multifunctional protein is primarily involved in genome stability maintenance through DNA damage repair. Thus, *BRCA1* mutation often results in erroneous DNA repair and cancer. Despite much work on BRCA1 protein’s recruitment to DNA damage sites and its role in multiple protein complexes, the direct interaction of BRCA1 protein with DNA is poorly understood. The evidence of BRCA1 protein’s direct interaction with DNA and strong affinity for HJ suggests its ability to recognize intermediates of DNA damage repair. However, the lack of structural information on the HJ-BRCA1 complex hampered our understanding of the molecular basis of BRCA1’s DNA recognition property. In our study, we have shown the structural preference of a full-length BRCA1 protein for an open X-like conformation over a stacked HJ for the first time at the singlemolecule level. The affinity of BRCA1 decreased an order of magnitude for stacked HJ. Generally, the junction binding proteins apparently prefer to bind stacked HJ and distort it to an open X-like structure. The observation of BRCA1 protein for an open X-like structure can provide new insight into protein-HJ interaction. Further, we modeled the DBD of BRCA1 in exon 11 and performed molecular docking and all-atom molecular dynamics simulation for spontaneous binding. We observed that mostly charged and polar amino acids are involved in the HJ-BRCA1 complex. The docking and all-atom MD simulation aligned with each other in reporting the interacting residues. This suggests the consequential role these amino acids play in stabilizing the BRCA1-HJ complex. Interestingly, many of these amino acids are reported to be the place for missense mutation. This may explain why mutations in these amino acids have catastrophic effects on DSB repair through HR. We anticipate that our study can provide insight into the molecular basis of BRCA1 protein-mediated DNA damage repair. With further biophysical study on BRCA1 protein and DNA interaction we expect new avenues will open for treatment of BRCA1 mutation.

## AUTHOR INFORMATION

### Author Contributions

SHK carried out the experiment and computational work, analyzed the data and wrote the manuscript. VK modeled the protein. NP analyzed the data and wrote and reviewed the manuscript.

## ACKNOWLEDGMENT

We thank IISER Tirupati for funding the work. SKH thanks IISER Tirupati for the fellowship.

We acknowledge the National Super Computing Mission (NSM) for providing computing resources for the HPC System, which is implemented by C-DAC and supported by the Ministry of Electronics and Information Technology (MeitY) and Department of Science and Technology (DST), Government of India.

