## Supporting Information for "Preferential binding of human BRAC1 protein with open X-like conformation of Holliday junction, a homologous recombination intermediate"

Table S1. I-TASSER Scores of the models with the highest confidence from each modeling.

|  | C-score | TM-score | RMSD |
| --- | --- | --- | --- |
| *Model1* | -3.72 | 0.31±0.10 | 14.4±3.7Å |
| *Model2* | -3.94 | 0.29±0.09 | 15.0±3.5Å |
| *Model3* | -3.94 | 0.29±0.09 | 15.0±3.5Å |

Table S2. Haddock scores of the docking of *Model1* (Docking 1), *Model2* (Docking 2) and *Model3* (Docking 3).

| Parameters | Docking 1 | | Docking 2 | | Docking 3 | |
| --- | --- | --- | --- | --- | --- | --- |
|  | Cluster 6 | Cluster 7 | Cluster 1 | Cluster 2 | Cluster 1 | Cluster 2 |
| HADDOCK score | 189.4 ± 31.2 | 211.1 ± 34.8 | 248.7 ± 10.3 | 265.0 ± 20.5 | 221.5 ± 39.2 | 221.8 ± 28.6 |
| Cluster size | 4 | 4 | 8 | 5 | 6 | 5 |
| RMSD from the overall lowest-energy structure | 1.9 ± 1.1 | 14.0 ± 0.2 | 13.5 ± 0.1 | 20.5 ± 0.4 | 11.7 ± 0.1 | 13.7 ± 0.1 |
| Van der Waals energy | -107.7 +/- 16.1 | -110.9 ± 25.7 | -97.5 ± 25.8 | -105.7 ± 7.1 | -113.1 ± 16.9 | -119.5 ± 8.0 |
| Electrostatic energy | -63.6 ± 72.2 | 49.6 ±79.3 | -84.0 ±125.2 | 11.1 ±52.0 | -155.9 ±83.0 | -152.0 ±33.9 |
| Desolvation energy | 23.3 ±5.5 | 27.3 ±3.2 | 34.5 ±9.8 | 17.7 ±1.9 | 35.2 ±5.6 | 34.9 ±4.1 |
| Restraints violation energy | 2879.6 ±167.86 | 2848.7 ±147.78 | 3456.0 ±264.33 | 3521.6 ±214.56 | 3429.8 ±304.82 | 3367.6 ±222.34 |
| Buried Surface Area | 2726.7 ±302.3 | 2780.7 ±500.3 | 2553.0 ±445.5 | 2456.3 ±126.6 | 2962.5 ±305.9 | 2767.0 ±140.3 |
| Z-Score | -1.6 | -0.9 | -1.2 | -0.7 | -1.1 | -1.1 |

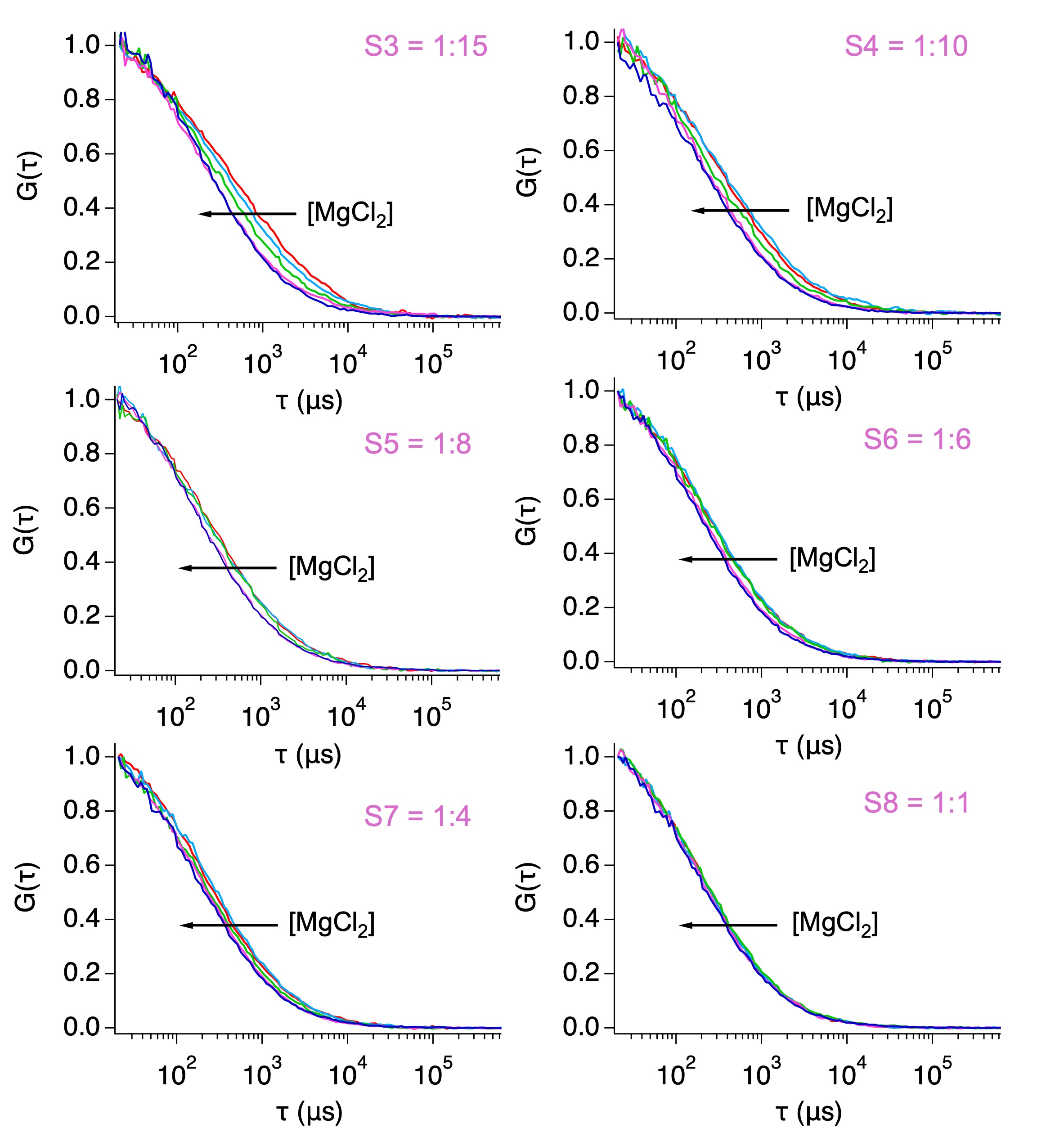

*Figure S1. Autocorrelation curves of HJ:BRCA1 = 1:15 (S3), 1:10 (S4), 1:8 (S5), 1:6 (S6), 1:4 (S4) and 1:1 (S8) with 0.5, 1.0, 5.0, 12.5 and 50 mM MgCl_2_.*

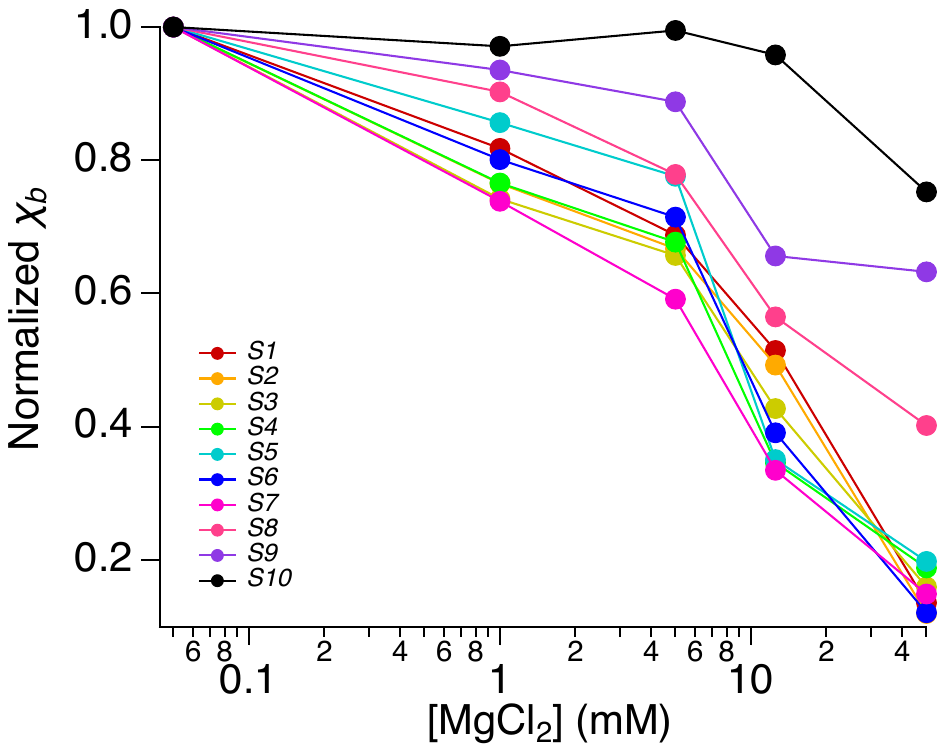

*Figure S2. Change in normalized* χ_b_ *with MgCl_2_.*

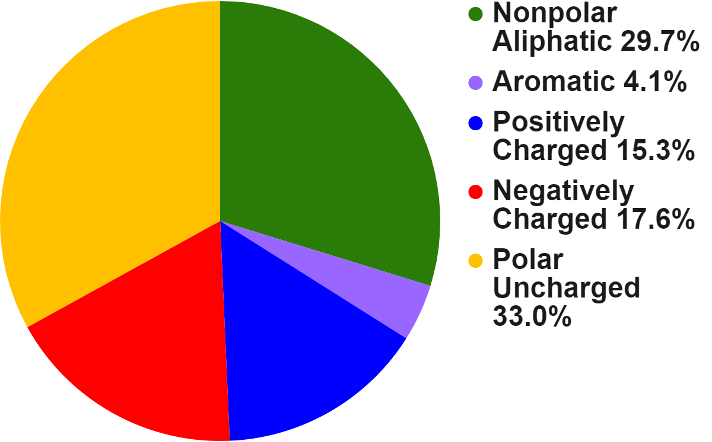

*Figure S3. Amino acid distribution of DBR (340-554) of BRCA1 protein.*

Table S3. vander Waal contacts found in docked complexes.

|  |  |  | No of VDW contacts | Nonpolar | Aromatic | Positively charged | Negatively charged | Polar uncharged |
| --- | --- | --- | --- | --- | --- | --- | --- | --- |
| Docking 1 | Cluster 6 | Complex32 | 115 | 11 | 11 | 44 | 27 | 22 |
|  |  | Complex86 | 101 | 14 | 9 | 22 | 21 | 35 |
|  |  | Complex180 | 120 | 19 | 8 | 36 | 36 | 21 |
|  |  | Complex188 | 138 | 20 | 9 | 38 | 37 | 34 |
|  | Cluster 7 | Complex71 | 164 | 12 | 24 | 37 | 48 | 43 |
|  |  | Complex76 | 76 | 8 | 6 | 10 | 29 | 23 |
|  |  | Complex85 | 134 | 25 | 12 | 37 | 29 | 31 |
|  |  | Complex167 | 165 | 26 | 6 | 39 | 39 | 55 |
|  | Total | | | 135 | 85 | 263 | 266 | 264 |
| Docking 2 | Cluster 1 | Complex14 | 141 | 12 |  | 60 | 37 | 32 |
|  |  | Complex30 | 87 | 8 |  | 61 | 10 | 8 |
|  |  | Complex32 | 150 | 23 |  | 60 | 39 | 28 |
|  |  | Complex82 | 144 | 22 |  | 67 | 29 | 26 |
|  | Cluster 2 | Complex18 | 122 | 30 |  | 46 | 31 | 15 |
|  |  | Complex23 | 126 | 44 |  | 36 | 12 | 34 |
|  |  | Complex102 | 145 | 40 |  | 50 | 23 | 32 |
|  |  | Complex137 | 141 | 43 |  | 53 | 16 | 29 |
|  | Total | | | 222 | 0 | 433 | 197 | 204 |
| Docking 3 | Cluster 1 | Complex15 | 141 | 17 | 2 | 33 | 22 | 67 |
|  |  | Complex39 | 178 | 14 | 1 | 62 | 19 | 82 |
|  |  | Complex55 | 127 | 22 | 1 | 38 | 15 | 51 |
|  |  | Complex137 | 94 | 10 |  | 18 | 11 | 55 |
|  | Cluster 2 | Complex8 | 175 | 27 |  | 62 | 37 | 49 |
|  |  | Complex51 | 138 | 20 | 1 | 57 | 26 | 34 |
|  |  | Complex89 | 166 | 34 | 1 | 46 | 27 | 58 |
|  |  | Complex91 | 126 | 22 |  | 33 | 37 | 34 |
|  | Total | | | 166 | 6 | 349 | 194 | 430 |

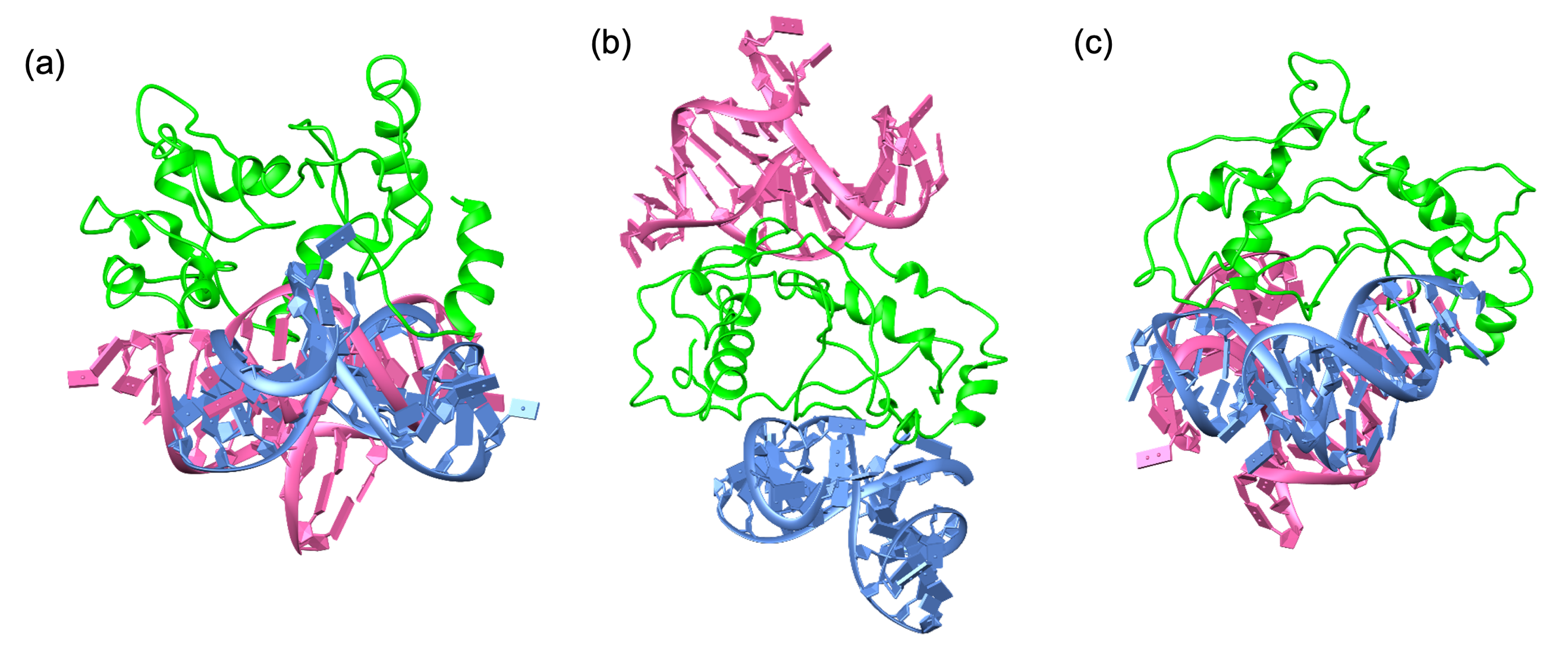

*Figure S4. Representative docked DBR-HJ complex of (a) Model1, (b) Model2 and (c) Model3. HJs in pink and light blue represent two docked structures from the best cluster in each docking.*

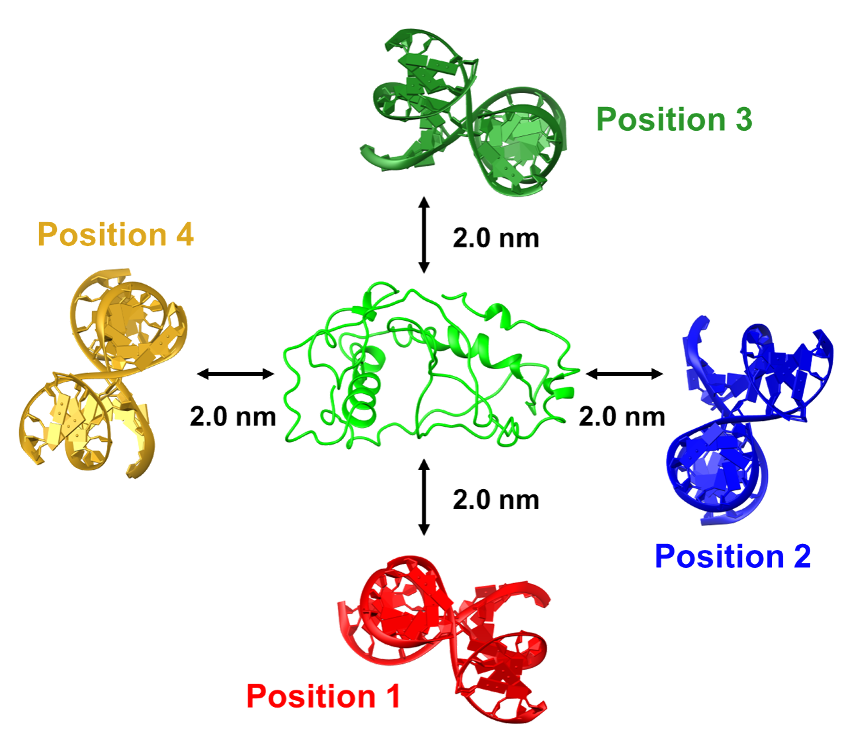

*Figure S5. Four starting orientations of HJ separated by a distance of ≈2 nm from model2 of DBR of BRCA1 to observe spontaneous binding in all-atom MD simulation.*

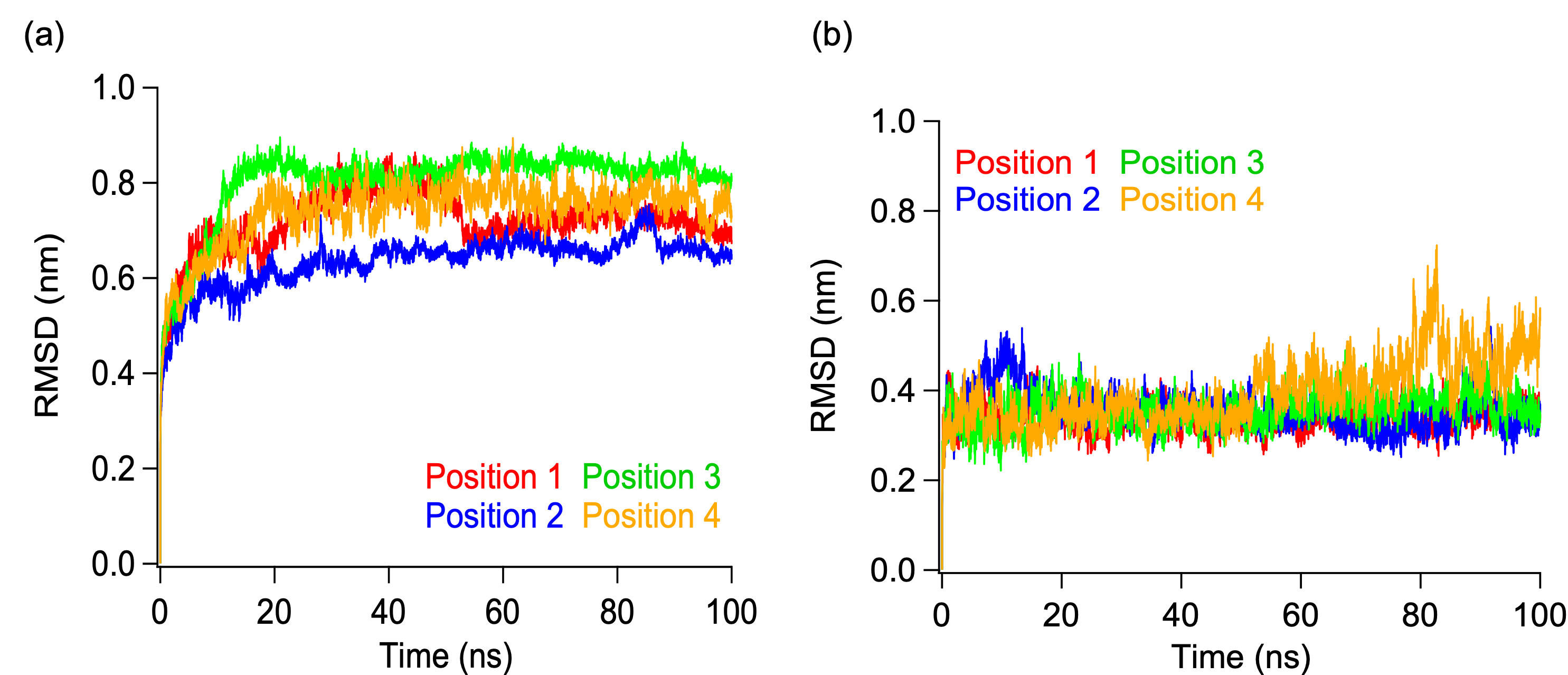

*Figure S6. RMSD plot of DBR of (a) BRCA1 and (b) HJ during spontaneous binding in all-atom MD simulation.*
